# Single-molecule diffusometry reveals no catalysis-induced diffusion enhancement of alkaline phosphatase as proposed by FCS experiments

**DOI:** 10.1101/2020.04.10.036442

**Authors:** Zhijie Chen, Alan Shaw, Hugh Wilson, Maxime Woringer, Xavier Darzacq, Susan Marqusee, Quan Wang, Carlos Bustamante

## Abstract

Theoretical and experimental observations that catalysis enhances the diffusion of enzymes have generated exciting implications about nanoscale energy flow, molecular chemotaxis and self-powered nanomachines. However, contradictory claims on the origin, magnitude, and consequence of this phenomenon continue to arise. Experimental observations of catalysis-enhanced enzyme diffusion, to date, have relied almost exclusively on fluorescence correlation spectroscopy (FCS), a technique that provides only indirect, ensemble-averaged measurements of diffusion behavior. Here, using an Anti-Brownian ELectrokinetic (ABEL) trap and in-solution spectroscopy (FCS), a technique that provides only indirect, ensemble-averaged measurements of diffusion behavior. Here, using an Anti-Brownian ELectrokinetic (ABEL) trap and in-solution single-particle tracking (SPT), we show that catalysis does not increase the diffusion of alkaline phosphatase (ALP) at the single-molecule level, in sharp contrast to the ~20% enhancement seen in parallel FCS experiments using *p*-nitrophenyl phosphate (*p*NPP) as substrate. Combining comprehensive FCS controls, ABEL trap, surface-based single-molecule fluorescence, and Monte-Carlo simulations, we establish that *p*NPP-induced dye blinking at the ~10 ms timescale is responsible for the apparent diffusion enhancement seen in FCS. Our observations urge a crucial revisit of various experimental findings and theoretical models––including those of our own––in the field, and indicate that in-solution SPT and ABEL trap are more reliable means to investigate diffusion phenomena at the nanoscale.

**SIGNIFICANCE STATEMENT:** Recent experiments have suggested that the energy released by a chemical reaction can propel its enzyme catalyst (for example, alkaline phosphatase, ALP). However, this topic remains controversial, partially due to the indirect and ensemble nature of existing measurements. Here, we used recently developed single-molecule approaches to monitor directly the motions of individual proteins in aqueous solution and find that single ALP enzymes do not diffuse faster under catalysis. Instead, we demonstrate that interactions between the fluorescent dye and the enzyme’s substrate can produce the signature of apparent diffusion enhancement in fluorescence correlation spectroscopy (FCS), the standard ensemble assay currently used to study enzyme diffusion and indicate that single-molecule approaches provide a more robust means to investigate diffusion at the nanoscale.

## INTRODUCTION

At the nanoscale, passive Brownian diffusion dominates the mobility of molecules. Whether freely-diffusing enzymes can harness chemical energy to generate additional mobility on top of Brownian motion is not well understood (1–3). Such a possibility seems to be supported by recent FCS measurements, which have shown that a number of non-motor enzymes including F1-ATPase (4), urease (5–8), catalase (6, 9), ALP (6), fructose bisphosphate aldolase (10), acetylcholinesterase (7) and hexokinase (11), enhance their diffusivities in the presence of substrates. Accordingly, various mechanisms including fluctuations in pH (5), global temperature increase of the solution (12), force and charged product generation (5), oligomeric enzyme dissociation (13), and enzyme chemotaxis toward substrates (9), have been proposed to account for this phenomenon. Using FCS, we previously proposed a mechanistic link between the enhanced diffusion of catalase, urease and ALP, and the heat released during their exothermic reactions (6). Within the framework of a stochastic theory, we proposed a “chemoacoustic effect” in which the heat released during catalytic turnover generates an asymmetric pressure wave that displaces the center-of-mass of the enzyme, manifesting as catalysis-enhanced enzyme diffusion. Arguing against this hypothesis, Illien *et al.* reported that aldolase, an enzyme that catalyzes a slow and endothermic reaction, also exhibits enhanced diffusion in the presence of its substrate or a competitive inhibitor (10), which they show to be independent of the overall turnover rate of the reaction. These authors propose that the enhanced diffusion is due to conformational fluctuations that alter the enzyme’s hydrodynamic radius. More surprisingly, a catalytically inert tracer has also been reported to diffuse faster in the presence of active enzymes (14), leading to the suggestion that the energy released during enzyme catalysis can be transferred to and harnessed by its environment. Directly contradicting the FCS results by Illien *et al.*, Zhang *et al.* and Günther *et al.* recently reported no diffusion enhancement of aldolase using dynamic light scattering (15) and pulsed field gradient nuclear magnetic resonance (16), respectively. On the other hand, theory suggests that the energy required to account for the experimentally observed diffusion enhancement far exceeds the chemical power released in enzymatic reactions (17). In addition, the change in the hydrodynamic radius of the enzyme needed to rationalize the observed diffusion enhancement is unlikely (18). To date, no unified theory has been proposed to rationalize these experimental observations, and publications in the field, either experimental or theoretical, rely almost exclusively on the validity of diffusion measurements made using FCS (1).

In FCS, time trace of light emitted while fluorescently labeled enzymes traverse a diffraction-limited confocal volume are recorded and analyzed in terms of the intensity autocorrelation function (19, 20). Because this autocorrelation is calculated over many molecules diffusing in and out of the focal volume, FCS only yields ensemble and time-averaged information. To extract the diffusion coefficient (D, μm^2^/s), an accurate fitting model is required, but not always available (21). Many factors other than diffusion contribute to the shape of the autocorrelation function, for example, dye photophysics (20), sample heterogeneity (e.g. mixture of species with different D values), geometry of the confocal volume (22) and optical aberrations (23). Failure to account for these factors could lead to erroneous interpretations of FCS data, as highlighted in a recent publication (1).

These confounding effects prompted us to reexamine catalysis-enhanced enzyme diffusion using single-molecule techniques and performing additional control experiments. Here, in addition to FCS, we use in-solution single-particle tracking (SPT) and ABEL trap based diffusometry to cross examine the diffusion of ALP. Our results reveal that the enzyme substrate (*p*NPP) affects the photophysics of the dye and introduces artifacts in the FCS measurements. This finding sets the stage necessary for future investigation of enzyme diffusion at the nanoscale.

## RESULTS

### Catalysis is neither sufficient nor necessary for the apparent diffusion enhancement of ALP in FCS

Among enzymes that have been reported to exhibit catalysis-enhanced diffusion, ALP shows the highest diffusion enhancement with its substrate *p*NPP (6) (Figure 1A), the magnitude of which is difficult to reconcile with theoretical predictions based on energetic coupling between the enzyme and its environment (18). We therefore sought to investigate the enhanced diffusion of ALP by performing additional control experiments in FCS (Figure 1B and 1C), and by using alternative methods to measure diffusion at the single-molecule level (Figure 1D). We purified commercial bovine intestinal ALP by size-exclusion chromatography (Figure S1A) and fluorescently labeled the enzyme with JF646 or Atto647N (Figure S1C). JF646 was chosen for its superior brightness and photostability (24), as shown in recent SPT and localization microscopy experiments (25). Using FCS, we recorded fluorescence transients of JF646-labeled ALP for 300 seconds, calculated the autocorrelation function G(τ) every 10 seconds, and fitted G(τ) with a simple model to estimate the diffusion coefficient (D) for each 10-second window (Figure 2A-C). The average D value obtained over the 300-second window is 45.6 ± 2.9 μm^2^/s (Figure 2C, data from 0 to 300s). We then added 2 mM *p*NPP substrate to the same solution during data acquisition. Interestingly, fluorescence intensity was quenched by ~50% immediately after substrate addition (even though the volume added is negligible) but recovered to 70% of the initial intensity within 150 seconds, remaining relatively stable thereafter (Figure 2A, right panel). Such *p*NPP-induced fluorescence quenching effect was also observed in bulk experiments with the free dye (Figure S2A). Analysis of 300 seconds of data obtained post *p*NPP addition gave a mean D value for the enzyme of 56.1 ± 4.7 μm^2^/s (Figure 2C, data after 300s), corresponding to a 23% apparent diffusion enhancement of ALP. Notably, the dispersion in extracted D values increases after *p*NPP addition (Figure 2C, red crosses). Consistent with previous findings (6), the diffusion enhancement decreases as *p*NPP concentration is lowered (Figure 2D, the first 4 groups with *p*NPP). Control experiments adding buffer did not give rise to any diffusion enhancement, indicating that the effect observed with *p*NPP was not due to perturbation of the experimental setup during sample addition (Figure 2D, buffer group). Labeling ALP with the previously used Atto647N dye yielded similar results (Figure S2B). In summary, these experiments confirmed previous FCS experiments reporting ALP-enhanced diffusion in the presence of *p*NPP.

**Figure 1.**
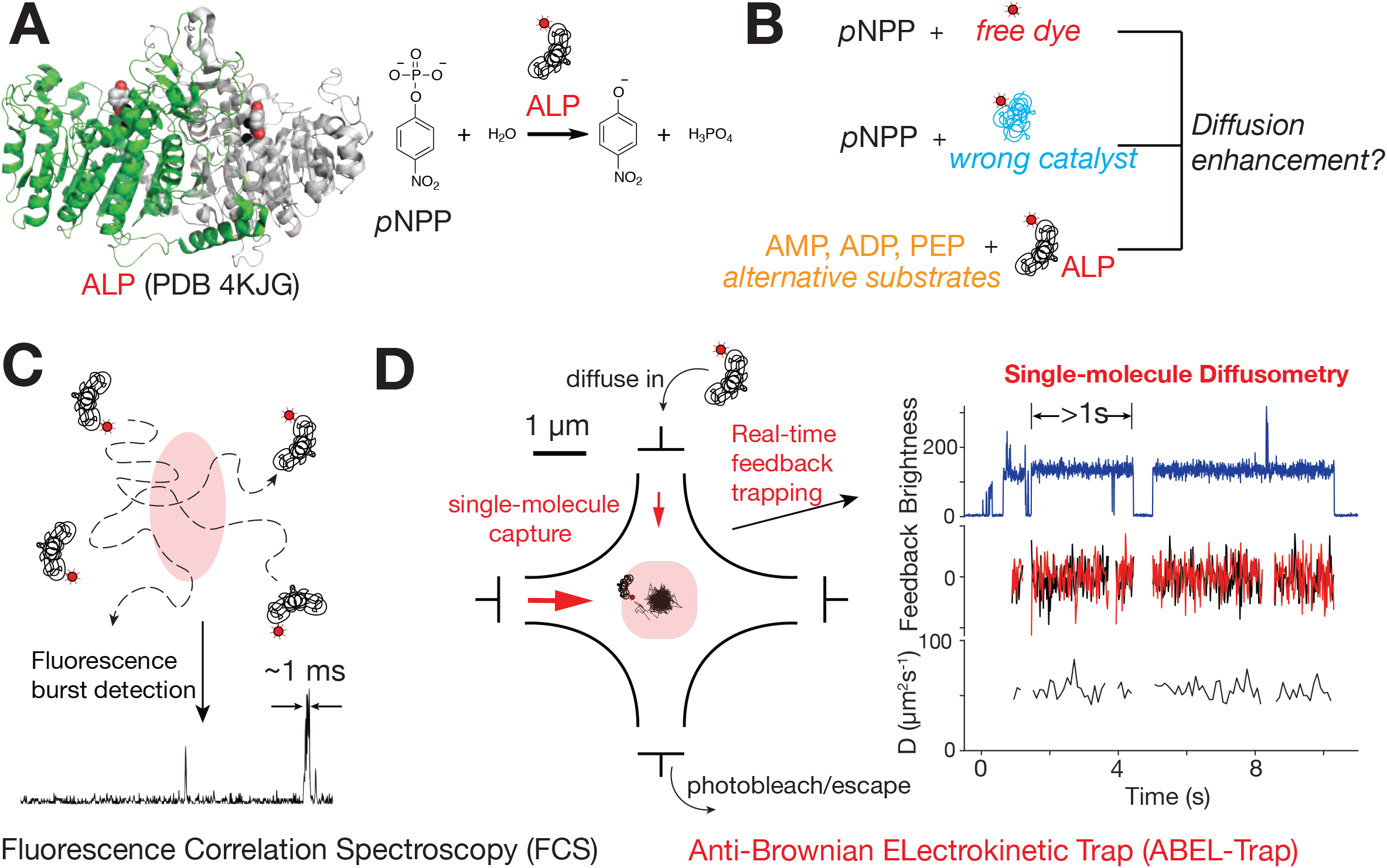
Revisiting Catalysis Enhanced Diffusion of ALP. (A) Structure of ALP (left panel) and the reaction it catalyzes (right panel). The two protomers of ALP are colored in green and grey. *p*NPP (red and white) and Zinc (black) are shown as spheres. During a reaction, ALP removes the phosphate group from *p*NPP and it has been shown previously that catalysis enhances the diffusivity of ALP. (B) Control experiments including *p*NPP with free dye, *p*NPP with wrong catalyst, and ALP reacting with alternative substrates are carried out to examine the role of catalysis on ALP diffusivity. (C) FCS estimates ensemble-averaged diffusion coefficients from autocorrelation analysis of fluorescence bursts in a confocal volume. (D) Principles of ABEL trap-based single-molecule diffusometry (see text for details). The diffusion coefficient of individual molecules can be measured for several seconds.

**Figure 2.**
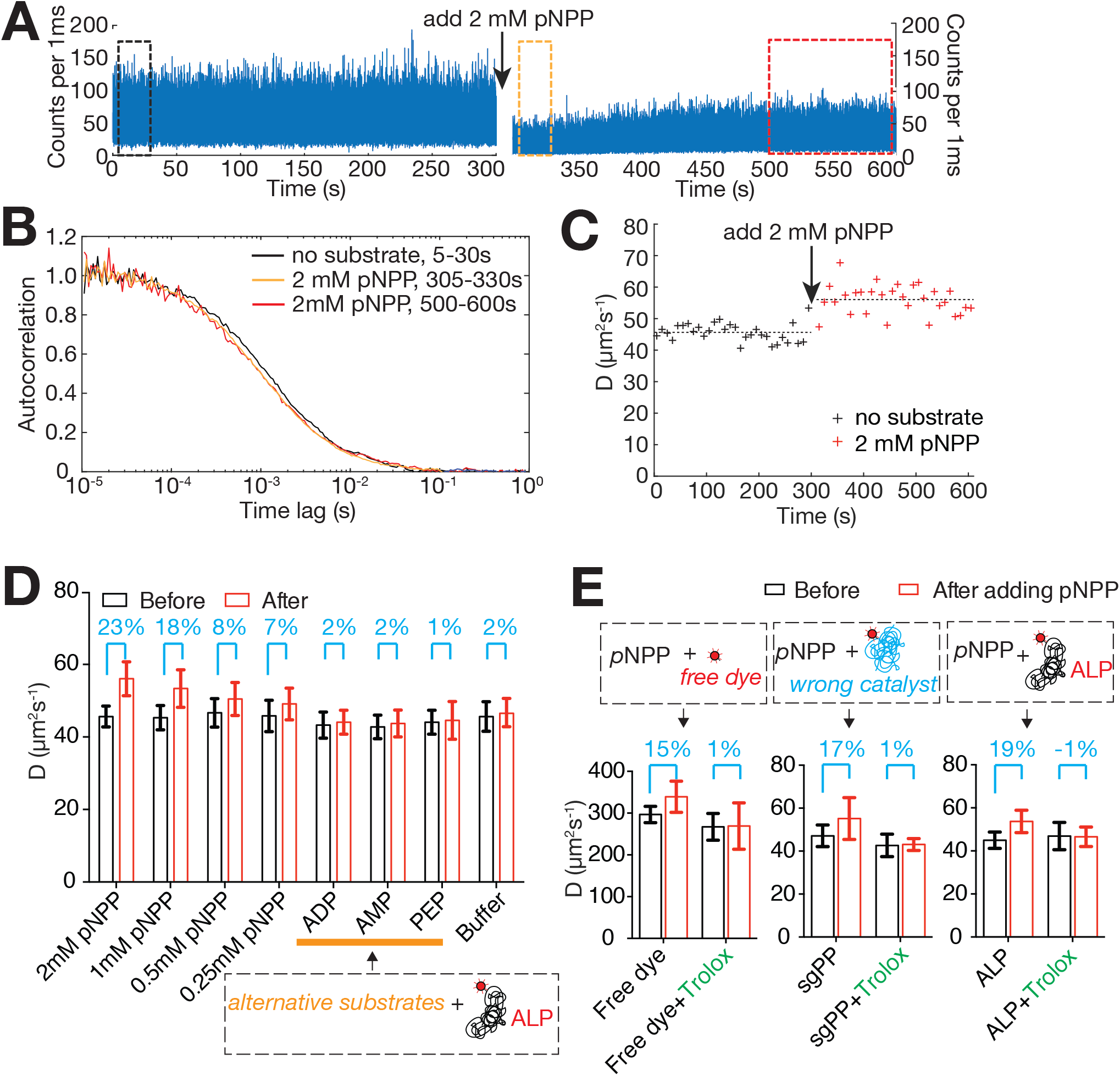
Catalysis is Neither Sufficient nor Necessary for Enhanced Diffusion of ALP in FCS. (A) FCS time trace of ALP-JF646 before (left) and after (right) adding 2 mM *p*NPP. (B) Normalized autocorrelation curves of data in black, orange and red rectangle boxes in (A). (C) D values extracted from fitting every 10s of data in (A). (D) *p*NPP, but not other substrates cause apparent diffusion enhancement of ALP. Mean D of 300s of data before (black bars) and after (red bars) adding the indicated solutions. Error bars are standard deviations of Ds extracted from 30 ten-second intervals. The percentage of mean D enhancement under each condition is labeled in cyan above the corresponding bars. The orange horizontal line indicates experiments with alternative substrates of ALP. (E) *p*NPP not only causes apparent diffusion enhancement of ALP, but also that of free dye and wrong catalyst. The apparent diffusion enhancement can be abrogated with a dye triplet state quencher, Trolox. The data is processed and represented similar to that of (D).

To test whether the diffusion enhancement of ALP observed in the presence of 2 mM *p*NPP originated from enzyme catalysis, we performed several control experiments. First, we measured free JF646 dye before and after addition of 2 mM *p*NPP. As shown in Figure 2E, addition of *p*NPP induced 15% apparent diffusion enhancement of free JF646 dye and a significant increase in the dispersion of D values (as characterized by the size of the error bars in that figure) (Figure 2E, free dye group), similar in magnitude to those seen in experiments with ALP-JF646. Second, we measured the diffusion coefficient of Atto647N-labeled *Streptococcus gordonii* inorganic pyrophosphatase (26) (*sg*PP-Atto647N, see supplementary methods for details) before and after addition of 2 mM *p*NPP. Even though *p*NPP is not a substrate of *sg*PP, we again observed a 17% apparent D enhancement and increase in the dispersion of D (Figure 2E, *sg*PP group). Third, we measured ALP-JF646 in the presence of other ALP substrates including 2 mM AMP (adenosine monophosphate), 4 mM ADP (adenosine diphosphate) and 2 mM PEP (phosphoenolpyruvate). ALP remain active with these substrates (Figure S1B) but no apparent D enhancement was observed with any of them (Figure 2D, ADP, AMP, PEP groups, and Figure S2B). Unlike *p*NPP, these compounds do not quench the fluorescence of JF646 dye in bulk (Figure S2A). Similar results were obtained when ALP was labeled with Atto647N dye. Collectively, these data indicate that catalysis is neither sufficient nor necessary for the apparent D enhancement of ALP in the presence of *p*NPP in FCS (Figure 2D-E).

### ABEL trap experiments reveal that *p*NPP induces dye quenching and blinking at the millisecond timescale

To further investigate the physical origin of the apparent diffusion enhancement observed in FCS, we turned to ABEL trap based single-molecule diffusometry (27), a recently developed technique capable of measuring the diffusion coefficient of individual molecules in solution. The ABEL trap senses the molecular position in real time via fluorescence detection and applies an electric voltage to exert an electrokinetic feedback control that counteracts the molecule’s Brownian motion (28). This procedure allows ~1-10 s continuous trapping of individual molecules—an observation time window three to four orders of magnitude longer than that in FCS (27, 29–31). The feedback voltages can then be used to reconstruct the diffusion trajectory and to estimate the diffusion coefficient of the individual molecules, directly. Unlike FCS, in which the emitted light intensity fluctuates due to both the number of molecules transiently traversing the confocal volume and the photophysics of the dye (e.g. quenching, blinking, etc), in the ABEL trap, the signal arises from a single molecule and variations in dye emission can be directly observed. Accordingly, the diffusion coefficient obtained in ABEL trap experiments are not subjected to artifacts arising from dye photophysics.

We first trapped single ALP-JF646 molecules without substrate. A typical dataset is shown in Figure 3A. Here, single molecules of ALP-JF646 diffuse into a ~3 μm × 3 μm trapping area and are captured for multiple seconds before escaping (usually due to photobleaching of the dye). After one molecule leaves, another one stochastically enters the area and becomes trapped, giving rise to a brightness trace (Figure 3A, top panel). Only one molecule is captured at a time and feedback voltages (Figure 3A, middle panel) are applied every time an emitted photon arrives at the detector to keep the molecule trapped. The feedback voltages are used to determine the average diffusion coefficient of the molecule (Figure 3A, bottom panel). Enzymes labeled with one or two JF646 dyes can be easily differentiated by their initial brightness and the number of discrete photobleaching steps (Figure 3A, top left panel). Due to the non-specific labeling scheme used here, we cannot draw any conclusion regarding the oligomerization state of the enzyme (i.e. monomer or dimer) from molecule brightness. On the other hand, the diffusion coefficient can be used to infer the oligomerization state of the protein (31). D values extracted from many single molecules are narrowly distributed around 54.8 μm^2^/s (Figure 3A, right histogram). This value matches well the diffusion coefficient (55.8 μm^2^/s) predicted from a hydrodynamic model (32) based on the enzyme’s crystal structure (PDB 4KJG), suggesting that the enzyme exists as a stable homodimer within our experimental time window (1-2 hours) and enzyme concentrations (20 pM).

**Figure 3.**
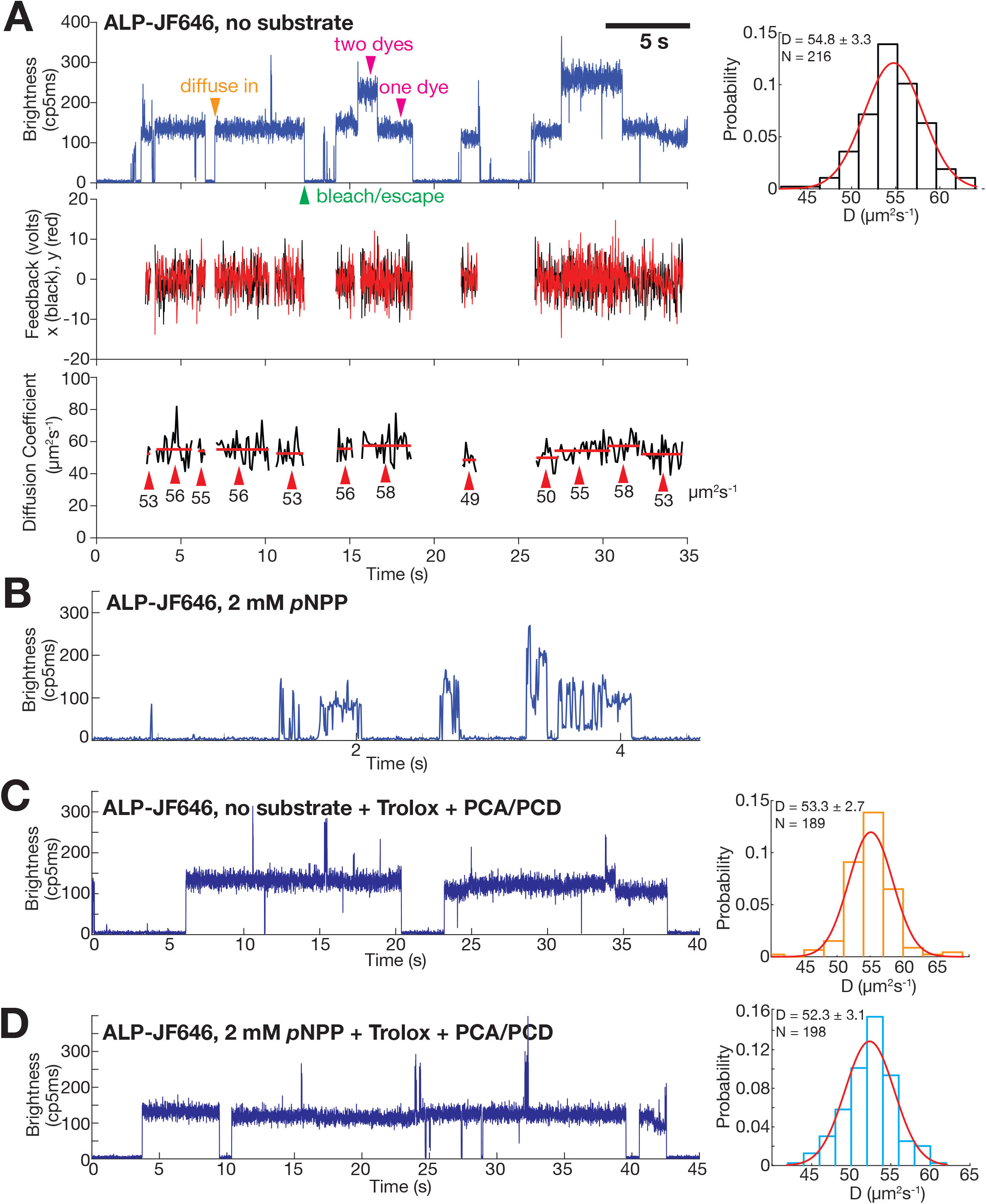
ABEL trap-based single-molecule diffusometry Reveals no Catalysis-induced Diffusion Enhancement of ALP. (A) A representative ABEL trap trace of ALP-JF646 with no substrate. Top left panel, brightness plot (fluorescence intensity, photon counts per 5 ms) of the detected fluorescence signal. A single fluorophore corresponds to a brightness level of ~130 (counts/5ms), whereas dual-fluorophore corresponds to ~260 (counts/5ms). Transient spikes in the brightness trace are caused by brief co-occupancy of two molecules in the trapping region. Orange arrow (diffuse in) denotes the start of a successful trapping event. Magenta arrows denote the regions of the data that corresponds to one or two dyes per protein molecule. Green arrow denotes the end of a trapping event, due to either photobleaching of the dye or escape of the trapped molecule. Middle panel, the corresponding feedback voltages (x in black, y in red) applied in order to counteract Brownian motion and keep the molecule in trap. The feedback is only plotted when there is a molecule in the trap. Bottom panel, D of each trapped molecule calculated every 100 ms (black trace). The red lines indicate the mean of each molecule, with the value written under the red lines and denoted with red arrows. The mean D of each molecule is used to build up the histogram on the top right panel. Top right panel, D histogram of ALP-JF646 without substrate and a Gaussian fit to the data. Mean ± Std of D and number of single molecules trapped (N) are displayed on the top left corner. (B) A representative ABEL trap intensity trace of ALP-JF646 with 2mM *p*NPP. (C) A representative ABEL trap intensity trace (left) and D histogram (right) of ALP-JF646 with no substrate in the Trolox + PCA/PCD buffer. (D) A representative ABEL trap intensity trace (left) and D histogram (right) of ALP-JF646 with 2 mM *p*NPP in the Trolox + PCA/PCD buffer.

Next, we attempted to trap ALP-JF646 under catalysis condition by adding 2 mM *p*NPP. Under these conditions we could no longer trap stably individual ALP-JF646 molecules. Instead, only brief transients with large brightness fluctuations and lasting from milliseconds to hundreds of milliseconds were observed (Figure 3B). Because ABEL trap relies on photon detection to counteract diffusion, trapping is not stable when the dye enters frequently or for a prolonged period of time a dark state. Therefore, to characterize this photophysical effects induced by *p*NPP, we measured emission from single JF646 dyes attached to surface immobilized DNA duplexes (Figure S3A). Without *p*NPP, stable dye emission lasting tens of seconds was observed (Figure S3B). Addition of 2 mM *p*NPP resulted in rapid on-off switching (blinking) and a dramatic reduction of the on-time of the fluorophore (Figure S3C), confirming the quenching and blinking effects induced by *p*NPP seen in the ABEL trap. These observations are also consistent with the quenching observed in FCS and bulk experiments (Figure 2A and S2A).

Single-molecule fluorophore quenching and blinking have been studied extensively (33), and several additives are known to rescue the molecules from the dark state. Indeed, when we included a commonly used anti-blinking reagent (Trolox with oxygen removal by protocatechuic acid, PCA, and protocatechuic deoxygenase, PCD) (34, 35) in our trapping buffer, the *p*NPP-induced dye blinking effect was completely suppressed, allowing us to trap single ALP-JF646 molecules in the presence of *p*NPP (Figure 3D, left panel) for multiple seconds in the ABEL trap. Thus, by suppressing *p*NPP-induced dye quenching and blinking, we identified the experimental conditions that allowed us to study the diffusion behavior of individual enzyme molecules under catalysis.

### ABEL trap reveals no catalysis-enhanced diffusion of ALP

Using this buffer system, we compared the distributions of D values obtained for ALP-JF646 with and without *p*NPP. In the presence of the anti-blinking reagent and in the absence of substrate, ALP molecules showed a narrow distribution (D = 53.3 ± 2.7 μm^2^/s, N = 189) (Figure 3C), similar to the values obtained in the absence of the reagent (Figure 3A). With 2 mM *p*NPP, we saw no difference in the distribution of the diffusion coefficient (D = 52.3 ± 3.1 μm^2^/s, N = 198) (Figure 3D). Notably, the enzyme remains active in all buffer conditions (Figure S1D). Thus, the ABEL trap data do not agree with the FCS results and instead indicate that there is no catalysis-induced diffusion enhancement of ALP.

### SPT reveals no catalysis-enhanced diffusion of ALP

To cross validate the new single-molecule observation that ALP does not diffuse faster under catalysis conditions, we performed high-speed, in-solution single particle tracking (SPT) experiments of the enzyme with and without *p*NPP. SPT has emerged as a powerful approach to track the movement of individual molecules (24, 36). By imaging fluorescent molecules at high rate, it is possible to localize their positions in successive frames (Figure 4A). The jump length distribution from these time courses can be used to derive their diffusion coefficients, D (Figure 4B) (37, 38). Unlike FCS, this method is less sensitive to dye photophysics, as dark state or blinking fluorophores will not appear in successive images for reconstruction of diffusion time courses. Using high-speed SPT with stroboscopic illumination (38) (Supplementary methods), we validated the robustness of this method (Figure S4) and imaged ALP-Atto647N at ~600 Hz (1.7 ms per frame) in buffer containing 10% glycerol, to slow down the diffusion. The acquired traces were then analyzed using a population model (37) that expresses the distribution of jump lengths assuming a Brownian, free-diffusion model (Figure 4B). Fitting both histograms of jump lengths to the model yielded similar diffusion coefficients with and without substrate (D = 22.5 μm^2^/s with no substrate, and D = 23.3 μm^2^/s with 5 mM *p*NPP, Figure 4C), confirming the ABEL trap finding that catalysis does not enhance the diffusivity of ALP at the single-molecule level. Consistent with the quenching and blinking effects of *p*NPP on the dye, we detected approximately 10-fold fewer particles in the presence than in the absence of *p*NPP (Figure S5A) and the intensity of the detected spots was also lower (Figure S5B).

**Figure 4.**
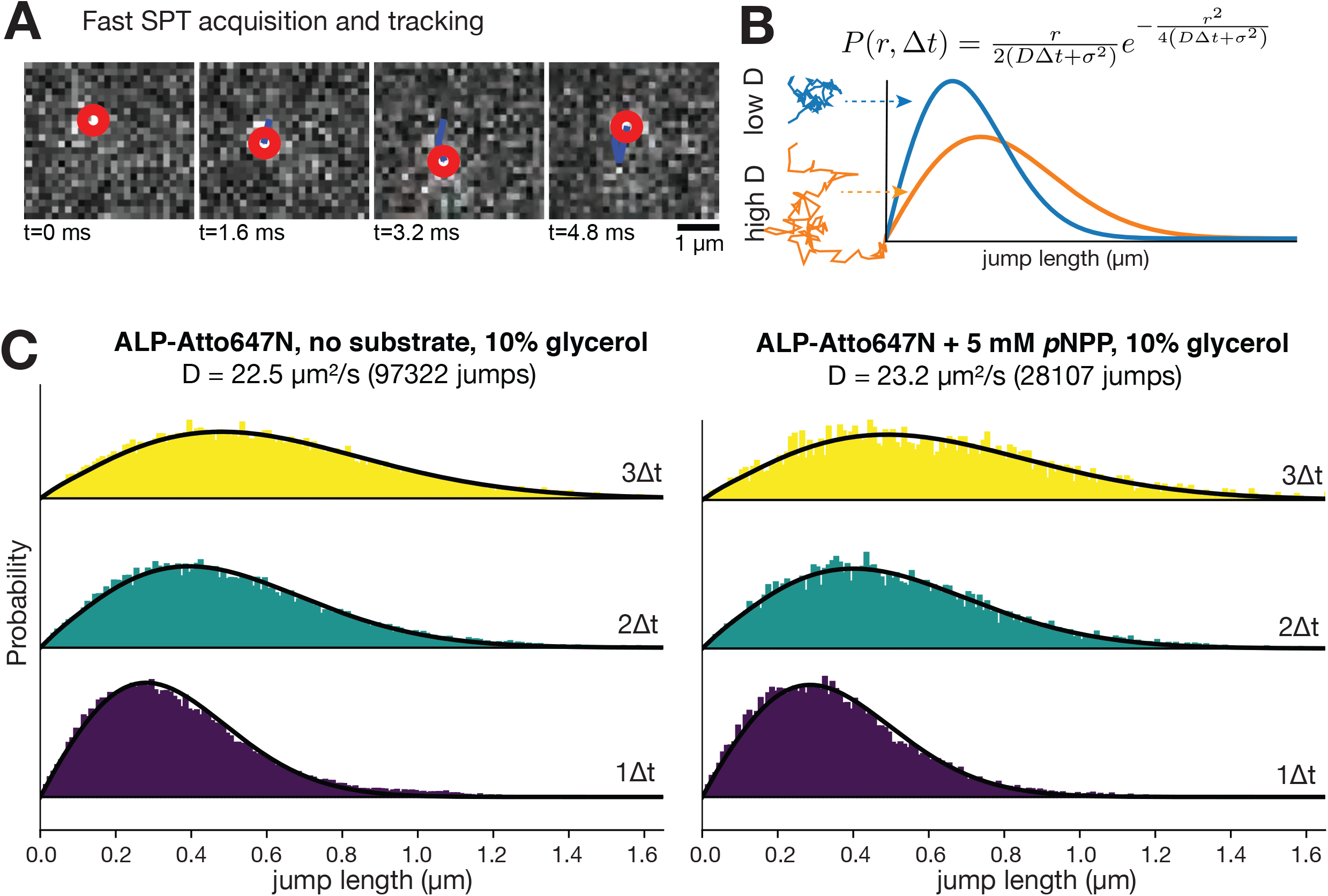
SPT Reveals no Catalysis-induced Diffusion Enhancement of ALP. (A) Representative images of high-speed SPT acquisition and tracking of single molecules. The red circle denotes the center of the detected molecule in each frame. The blue line denotes the molecule’s diffusion trajectory. (B) Modeling jump length distribution under free diffusion assumptions. Jump length distribution histograms of molecules with low and high Ds are plotted in blue and orange, respectively. (C) Jump length distribution histograms of ALP-Atto647N with no substrate (left) and with 5 mM *p*NPP (right). The yellow, green and purple histograms are distributions of jump lengths between 3Δt (Δt =1.7 ms), 2Δt and 1Δt, respectively. The calculated D and the number of total jumps are written on the top of the histograms.

### The apparent diffusion enhancement of ALP in FCS is caused by *p*NPP-induced photophysics of the dye

The new single-molecule results prompted us to reexamine our previous FCS experiments. Because FCS relies on fitting the decay of the intensity correlation function in the 0.1-50 ms range to extract the diffusion coefficient, we hypothesize that *p*NPP-induced dye blinking––which was observed at ~10 ms timescale in the ABEL trap experiments––is responsible for the apparent diffusion enhancement of ALP in FCS. To test this hypothesis, we carried out FCS experiments in Trolox-based antiblinking buffer, which suppresses *p*NPP-induced dye blinking (Figure 3D). Significantly, no *p*NPP-induced diffusion enhancement of ALP-JF646, free JF646 dye and *sg*PP-Atto647N was observed (Figure 2E, + Trolox groups), confirming the hypothesis that *p*NPP-induced dye photophysics is responsible for the apparent diffusion enhancement of ALP in FCS.

By performing ABEL trap experiments of ALP-JF646 under different buffer conditions, we were able to elucidate the mechanism of *p*NPP-induced dye blinking. First, we found that not all components in the Trolox-based antiblinking mixture are needed to suppress blinking in the presence of 2 mM *p*NPP. A single component, 2 mM PCA, is sufficient (Figure S6B). However, 2 mM PCA does not prevent blinking of the dye (Figure S6A) in the presence of sub-stoichiometric quantities of *p*NPP (200 μM, O_2_ removed), suggesting that matching concentrations of *p*NPP and PCA are needed for stable emission. All of our observations regarding *p*NPP-induced JF646 photophysics (Figure 5A) can be reconciled by the reducing and oxidizing mechanism of dye quenching proposed by Vogelsang *et al.* (39). To achieve a stable emission in the redox framework, both a reducing and an oxidizing agent are needed in the absence of oxygen to efficiently depopulate the triplet state (T_1_), as well as the radical cation (F*+) or anion states (F*−), which result from the oxidation and reduction of the triplet state, respectively. Here, *p*NPP contains a nitro group, making it a plausible oxidizing agent; PCA is a known reducing agent (40) and Trolox, when dissolved in solution, contains both reducing and oxidizing components (41). We thus propose that *p*NPP quenches JF646 fluorescence by random collision and subsequent oxidization of JF646 triplet state to create a long-lived, cation state (F*+) (Figure 5B). If this cation state is not efficiently scavenged by a reductant (e.g. PCA and/or Trolox) in solution, the fluorophore stays dark for extended periods (>10 ms). This mechanism is quite general and suggests that *p*NPP-induced quenching is not limited to the JF646 dye. The same mechanism has been shown to operate with other red dyes of different structural families (39). Indeed, when we measured the emission of single immobilized DNA duplexes labeled with the Atto647 dye (Figure S7A and S7B), addition of 2 mM *p*NPP (Figure S7C) induces rapid blinking of the dye on the timescale of ~10 ms (Figure S7C). On the other hand, addition of 2.5 mM PEP (a redox-inactive substrate of ALP) does not induce blinking (Figure S7D). Recently, Günther et al. reported quenching of Alexa 488 by *p*NPP, manifested as a reduction of fluorescent lifetime of the dye in the presence of 2-4 mM of this substrate (1).

**Figure 5.**
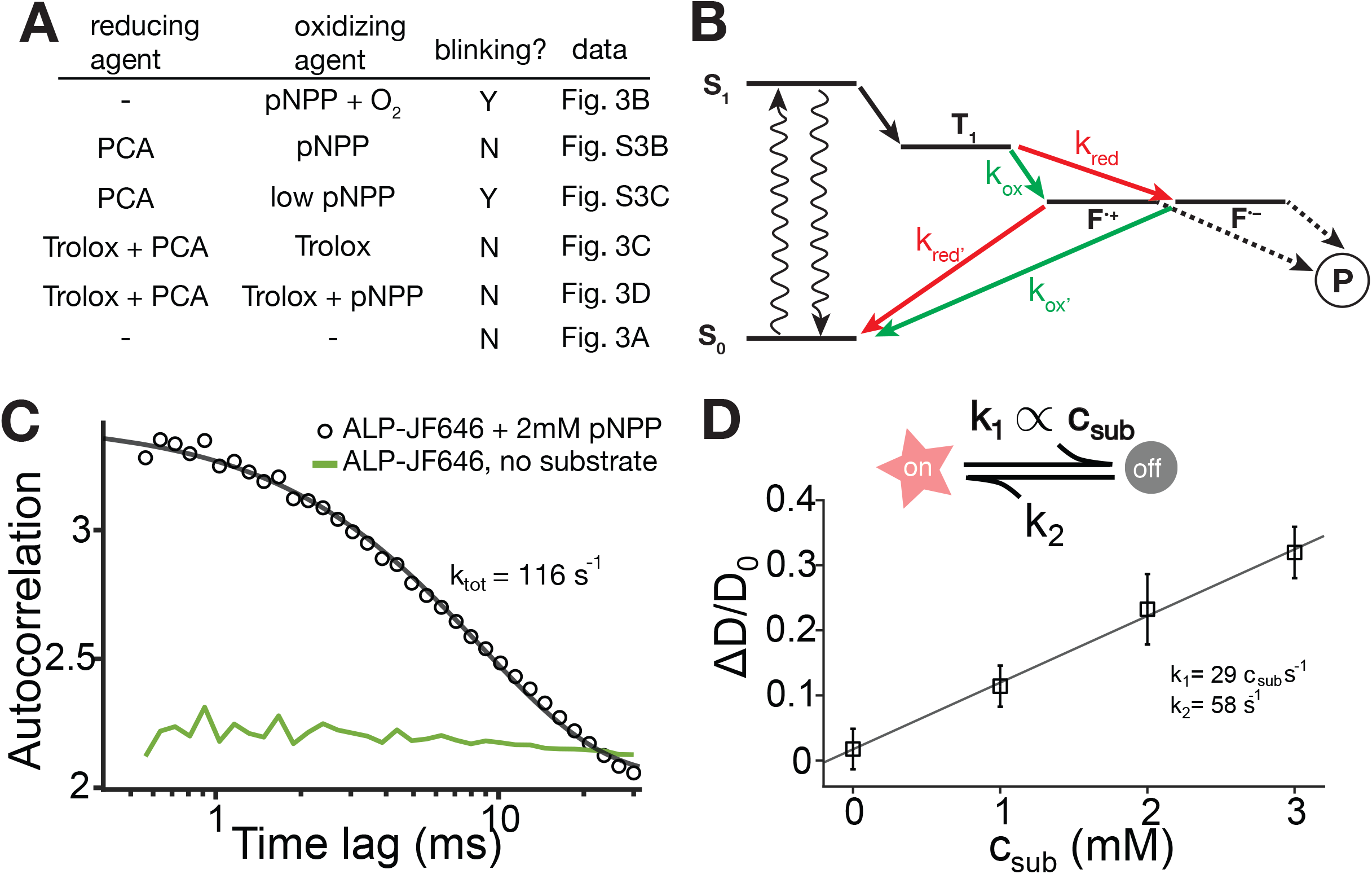
*p*NPP-induced Dye Photophysics is Responsible for the Apparent Diffusion Enhancement of ALP in FCS. (A) Summary of all ABEL trap ALP-JF646 experiment buffer conditions with identified oxidizing and reducing agents and blinking outcome, showing consistency with the reducing and oxidizing framework as depicted in (B). (B) A proposed model of *p*NPP-induced dye photophysics based on the reducing and oxidizing framework. The dye triple state (T_1_) can be oxidized (green arrows) or reduced (red arrows) into charge separated states (F^•+^ and F^•−^) which are not fluorescent and maybe prone to photobleaching (P). An oxidizing agent in solution needs to be balanced by the presence of a reducing agent (and vice versa) for blinking suppression, otherwise, the molecule is trapped in a dark radical state (F^•+^ or F^•−^) and blinks off. (C) Extracting *p*NPP-induced blinking kinetics from ABEL trap data. Intensity autocorrelation curve of ALP-JF646 with 2 mM *p*NPP (black circles, from the data represented by Figure 3B) shows a pronounced decay between 1ms and 30ms and is fitted with a single exponential function (black line) to extract the total rate. The intensity autocorrelation of ALP-JF646 without substrate (from Figure 3A) is shown in green for reference (amplitude multiplied by 10 for clarity). (D) Monte-Carlo simulation of FCS experiments with a substrate-induced blinking model (top schematic, see text and Supplementary for details), using the listed parameters constrained by the rate extracted in (C). The extracted apparent diffusion coefficient (averaged over 10 independent simulation runs) is plotted against the substrate concentration and fit with a line. Error bars represent standard deviation.

Finally, to evaluate quantitatively whether *p*NPP-induced dye photophysics can give rise to the magnitude (~20%) of apparent D enhancement seen in the FCS experiments, we conducted Monte-Carlo simulations of our FCS experiments (see supplementary text and Figure S8 for details), where the dye can switch from an “on” state to an “off” state with a rate (*k*_*1*_) that depends linearly on substrate concentration and a constant off-to-on rate (*k*_*2*_) that corresponds to spontaneous return of the dye to the ground state, S_0_ (Figure 5D). The total switching rate (*k*_*tot*_ = *k*_*1*_+*k*_*2*_ =116 s^−1^) at 2 mM *p*NPP was directly extracted from ABEL trap data (Figure 3B) by fitting the observed intensity autocorrelation function in the range of 0.5-20 ms to a single exponential decay (Figure 5C). Given that it is difficult to determine *k*_*1*_ and *k*_*2*_ uniquely in our experiments, we used *k*_*1*_ = *k*_*2*_ = *k*_tot_/2 as a rough estimate. The simulated intensity trace was subjected to autocorrelation analysis and fitted with the same one-species model without photophysics (Figure S8). The simulation results successfully recapitulate the magnitude (~20%) of the apparent D enhancement seen in our FCS experiments (Figure 5D).

Taken together, all the results presented above indicate that *p*NPP-induced dye blinking is responsible for the apparent diffusion enhancement of JF646, Atto647N, ALP-Atto647N and ALP-JF646 observed in FCS.

## DISCUSSION

Catalysis-fueled propulsion of biomolecules at the nanoscale is no doubt a fascinating concept and has potential applications in the field of nanoscience and medicine (42–49). However, mounting experimental and theoretical evidence argue against the mechanism, scale, and even the existence of such phenomenon (1, 2, 15, 17, 18, 50). The vast majority of publications documenting enhanced enzyme diffusion upon catalysis were performed with FCS, which is prone to artifacts, such as free dye contamination, dissociation of enzyme quaternary structure and dye photophysics, that could result in apparent enhanced enzyme diffusion. A deeper experimental and theoretical investigation of this phenomenon calls for better tools, and direct single-molecule measurements of molecular diffusion at the nanoscale could offer the clearest picture.

Here, we have taken the first steps towards this goal. By directly measuring the diffusion coefficient of individual enzyme molecules, we uncover an apparent discrepancy between the diffusion behavior of ALP in the presence of *p*NPP measured with FCS (~20% enhancement) and with single-molecule techniques (no enhancement). On the basis of several crucial control FCS experiments, single-molecule diffusometry, dye photophysics measurements on the surface, and simulation, we conclude that the apparent diffusion enhancement of ALP observed in FCS is caused by *p*NPP-induced quenching and blinking of the dye. We propose that *p*NPP-induces dye quenching and blinking through a redox-based electron transfer mechanism. Notably, dye photophysics, depending on the mechanism, can have timescales ranging from nanoseconds to seconds. However, substrate-induced redox blinking, as proposed here, is particularly detrimental to diffusion estimates in FCS because its timescale (~10 ms) may coincide with the diffusion transit timescale of molecules through the confocal volume. For this reason, we urge caution in interpreting FCS measurements when adding compounds that have distinct redox properties (e.g. hydrogen peroxide).

Given the results presented here and similar concerns voiced by others (1), we believe that the experimental evidence for enhanced enzyme diffusion needs to be critically reevaluated. We advocate the use of single-molecule techniques such as in-solution SPT and ABEL trap based diffusometry. These methods avoid the potential artifacts of ensemble methods such as FCS and yield a full distribution of diffusion coefficients in a molecular population. This latter capability will allow us to test alternative hypotheses of enhanced diffusion, such as redistribution of enzyme oligomerization states associated with substrate binding or catalysis, which has been suggested for aldolase (15, 51). As examples of this new measurement capability, single-molecule diffusometry (by ABEL trapping) recently revealed the nucleotide-dependent shift of quaternary structure of rubisco activase between monomer, dimer and hexamers (31). Here we have confirmed that ALP molecules remain as dimers under our experimental conditions.

## Materials and Methods

A detailed description of the materials and methods is given in the Supplementary Information. Briefly, Alkaline phosphatase (ALP) from bovine intestinal mucosa was purchased from Sigma-Aldrich (Cat # P7923), further purified in house, and fluorescently labeled using amine-NHS chemistry. ABEL trap based single-molecule diffusometry was implemented as previously described (27) and high-speed SPT was carried out according to the recent protocol (37).

## ACKNOWLEDGEMENT

We thank Philip Tinnefeld for comments and suggestions of the manuscript. We are grateful to members in the Marqusee lab, Bustamante Lab, Darzacq lab and the Zimmer lab for insightful comments and discussions, to Ana Robles and Astou Tangara for microscope maintenance and setup, to Ronen Gabizon for experimental assistance and discussions. QW acknowledges Evangelos Gatzogiannis for the loan of a PI P-563 piezo stage. AS is supported by the Swedish Research Council International Postdoc Grant (2017-00389). This work was funded by a grant from the US Department of Energy, Office of Basic Energy Sciences, Division of Materials Sciences and Engineering under contract No.DE-AC02-05CH11231 (CB), the National Institutes of Health grant R01GM032543 (CB) and UO1-497 EB021236 (XD), the National Science Foundation grant MCB 1616591 (SM), and a Lewis-Sigler Fellowship of Princeton University (QW). SM is a Chan Zuckerberg Biohub Investigator. CB is an investigator with the Howard Hughes Medical Institute.

## FOOTNOTES

To whom correspondence may be addressed: S.M., Q.W. or C.B.

### Author contributions

Z.C., H.W., A.S., and Q.W. designed and performed FCS experiments; H.D., M.W., Z.C., and A.S. designed and performed SPT experiments; Q.W., Z.C., and H.W. designed and performed ABEL trap experiments; Q.W. performed simulation experiments; Z.C. and Q.W. designed and performed surface experiments of dye photophysics; Z.C., and A.S. purified and labeled the enzymes; A.S., and H.W. performed bulk fluorescence and enzyme activity measurements; Z.C., H.W., A.S., and Q.W. analyzed FCS data; M.W. analyzed SPT data; Q.W., Z.C., and H.W. analyzed ABEL trap data; Z.C. wrote the initial draft of the manuscript; all authors commented, reviewed and revised the manuscript. S.M., Q.W., and C.B. co-supervised the study.

The authors declare no conflict of interest.

---

**Figure S1.**
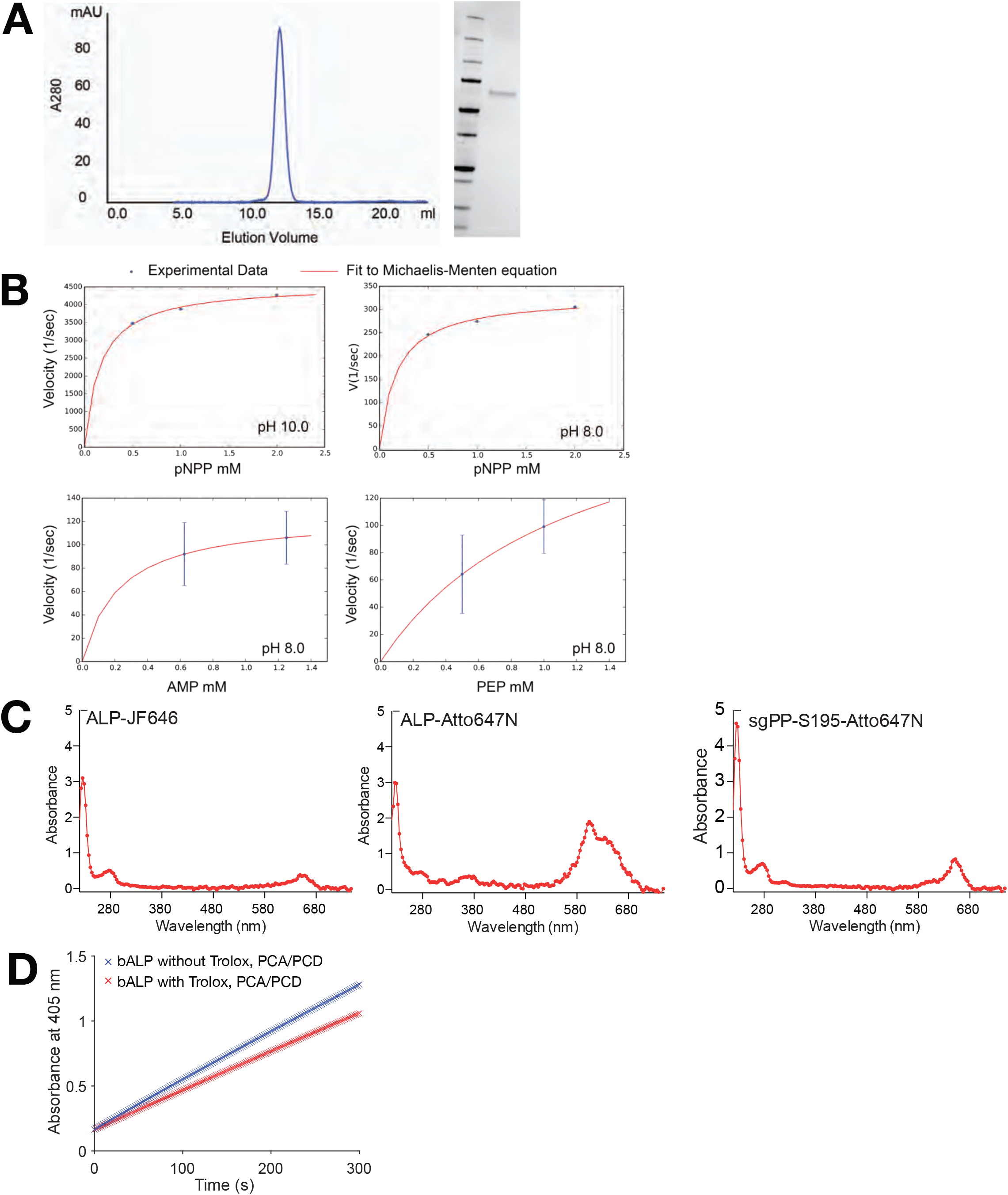
Purification, Labeling and Activity of ALP. (A) Size exclusion chromatography (left) and SDS-PAGE gel (right) of purified ALP. (B) Michaelis-Menten curves of ALP under different substrate and buffer conditions. (C) UV-Vis absorbance profiles of ALP-JF646, ALP-Atto647N and sgPP-S195C-Atto647N. (D) ALP is active in the presence and absence of Trolox and PCA/PCD. Absorbance of the reaction product 4-nitrophenol at 405 nm was monitored at room temperature for 5 minutes.

**Figure S2.**
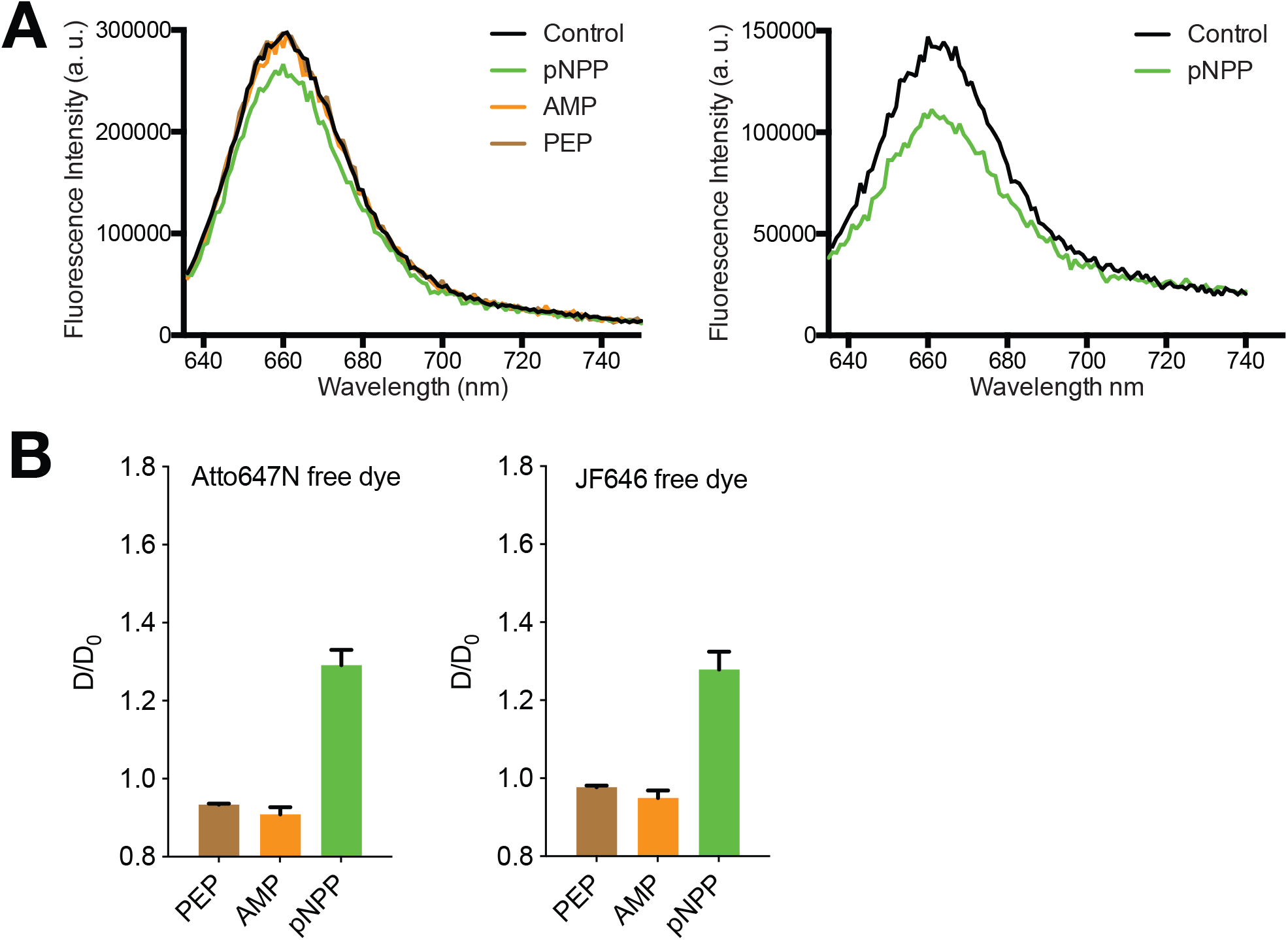
Effects of ALP Substrates on the Brightness and Diffusion Coefficient of Free Dyes. (A) *p*NPP quenches the fluorescence of Atto647N (left) and JF646 (right) as measured with a fluorometer. (B) *p*NPP, but not PEP or AMP causes apparent D enhancement (plotted as ratio of D/D_0_) of both free dye and ALP-dye in FCS. Dye is Atto647N and JF646 in the left and right panel, respectively.

**Figure S3.**
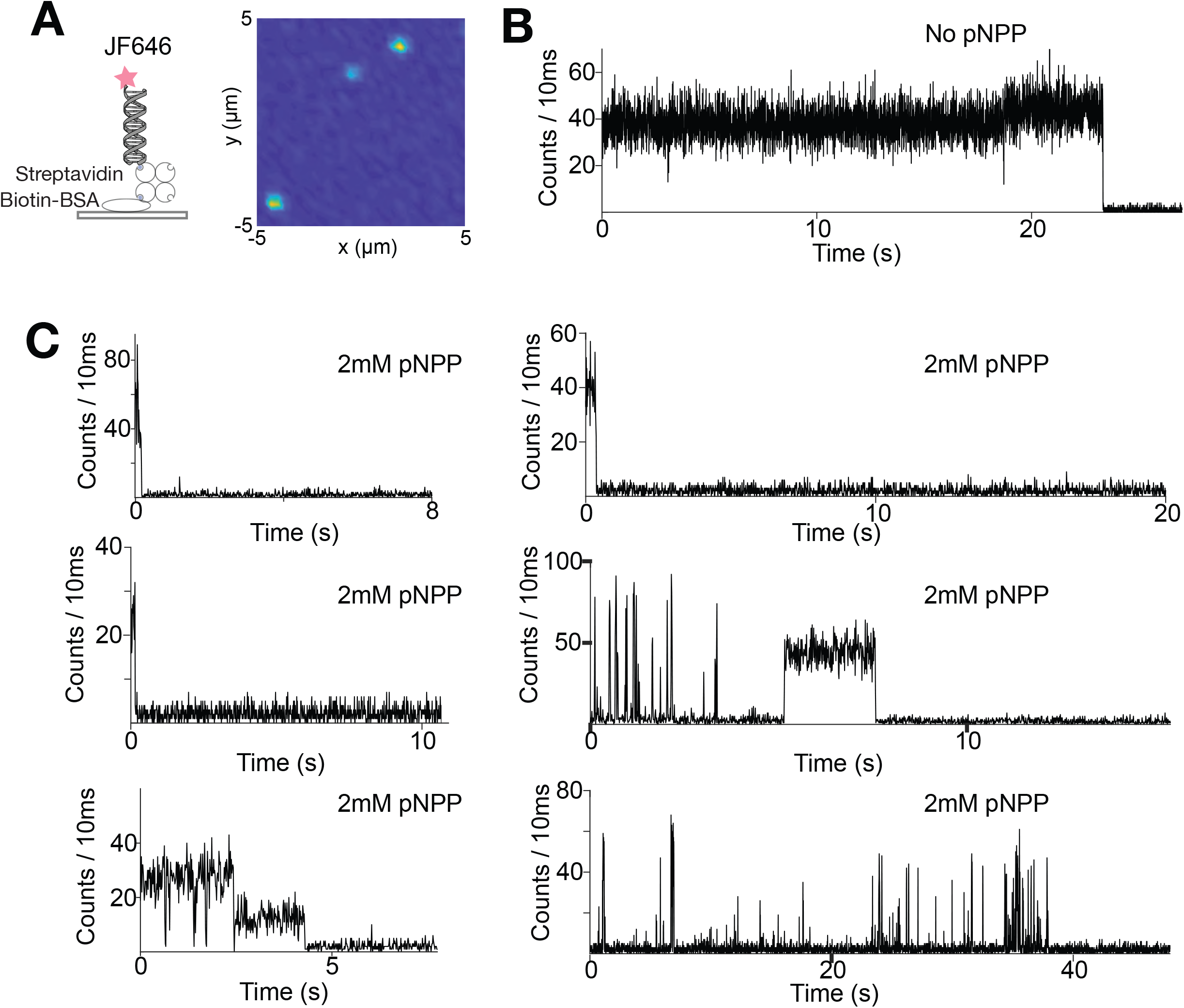
Surface-based single-molecule spectroscopy on JF646-labeled double-stranded DNA molecules. (A) Left, single JF646-dsDNA molecules are tethered to the surface via biotin-streptavidin interactions. Right, a representative image showing a 10μm ×10μm field of view, acquired ina buffer containing 20mM HEPES, 100mM NaCl without substrate. (B) A representative intensity (photon counts per 10 ms) trace of a single JF646-dsDNA molecule without *p*NPP. (C) Representative intensity traces of single JF646-dsDNA molecules in the presence of 2 mM *p*NPP. The “on” time of the molecules are drastically shortened and about 20% of the molecules show rapid blinking.

**Figure S4.**
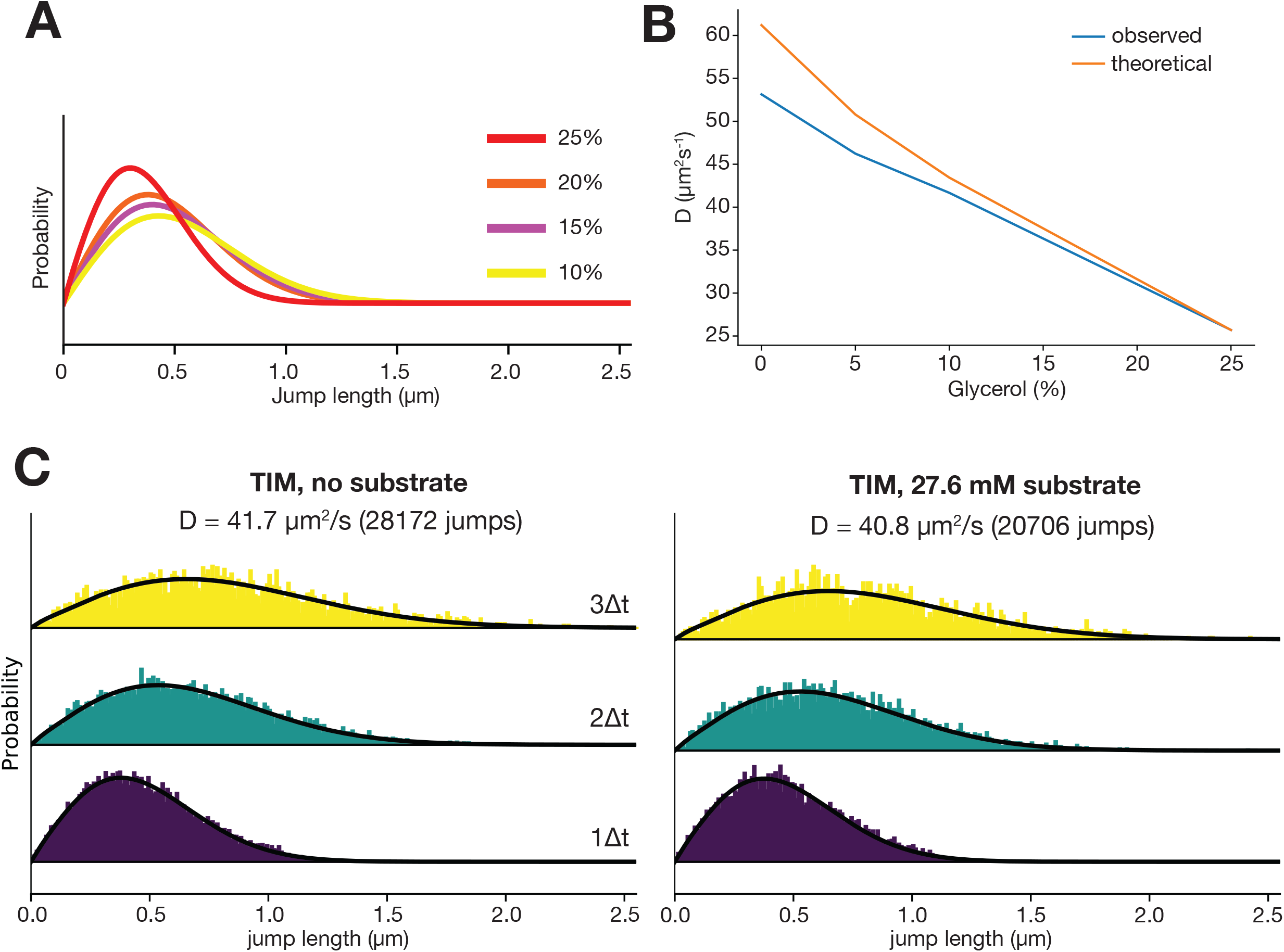
SPT control experiments. (A) Jump length distributions of TIM-Atto647 in buffers with different glycerol concentrations. (B) Measured D of TIM-Atto647 as a function of glycerol percentage and comparison to predicted D values using known viscosities of glycerol-water mixtures. (C) Jump length distribution histograms of TIM-Atto647 with no substrate (left) and with 27.6 mM of substrate D-glyceraldehyde 3 phosphate (right). The yellow, green and purple histograms are distributions of jump lengths between 3Δt, 2Δt and Δt (1.7 ms), respectively. The calculated D and the total number of jumps are shown on the top of the histograms.

**Figure S5.**
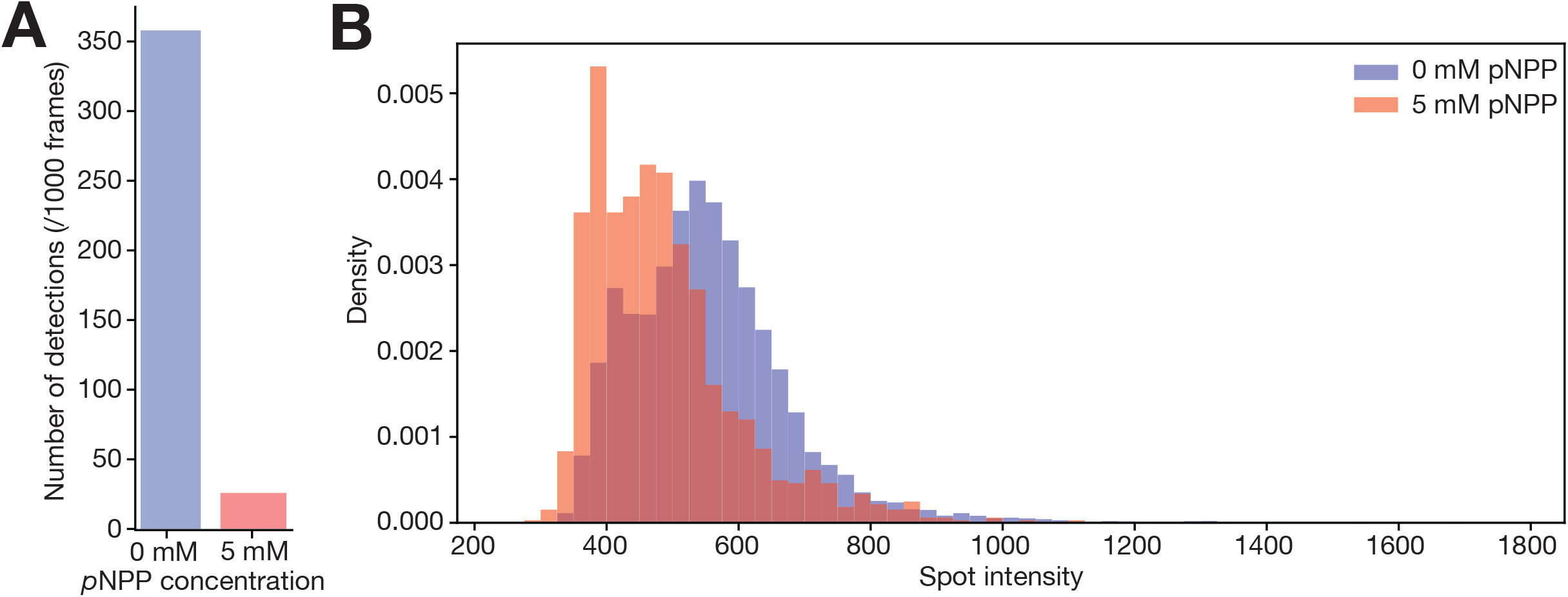
*p*NPP quenches dye fluorescence in SPT. (A) Number of detected particles (per 1000 frames) of ALP-Atto647N without (blue) and with 5 mM *p*NPP (orange) in SPT measurements. Much fewer particles are detected in the presence of *p*NPP due to quenching or blinking of the fluorophore. (B) Histograms of detected spot intensities without (blue) and with 5 mM *p*NPP (orange). The spot intensities are lower in the presence of *p*NPP.

**Figure S6.**
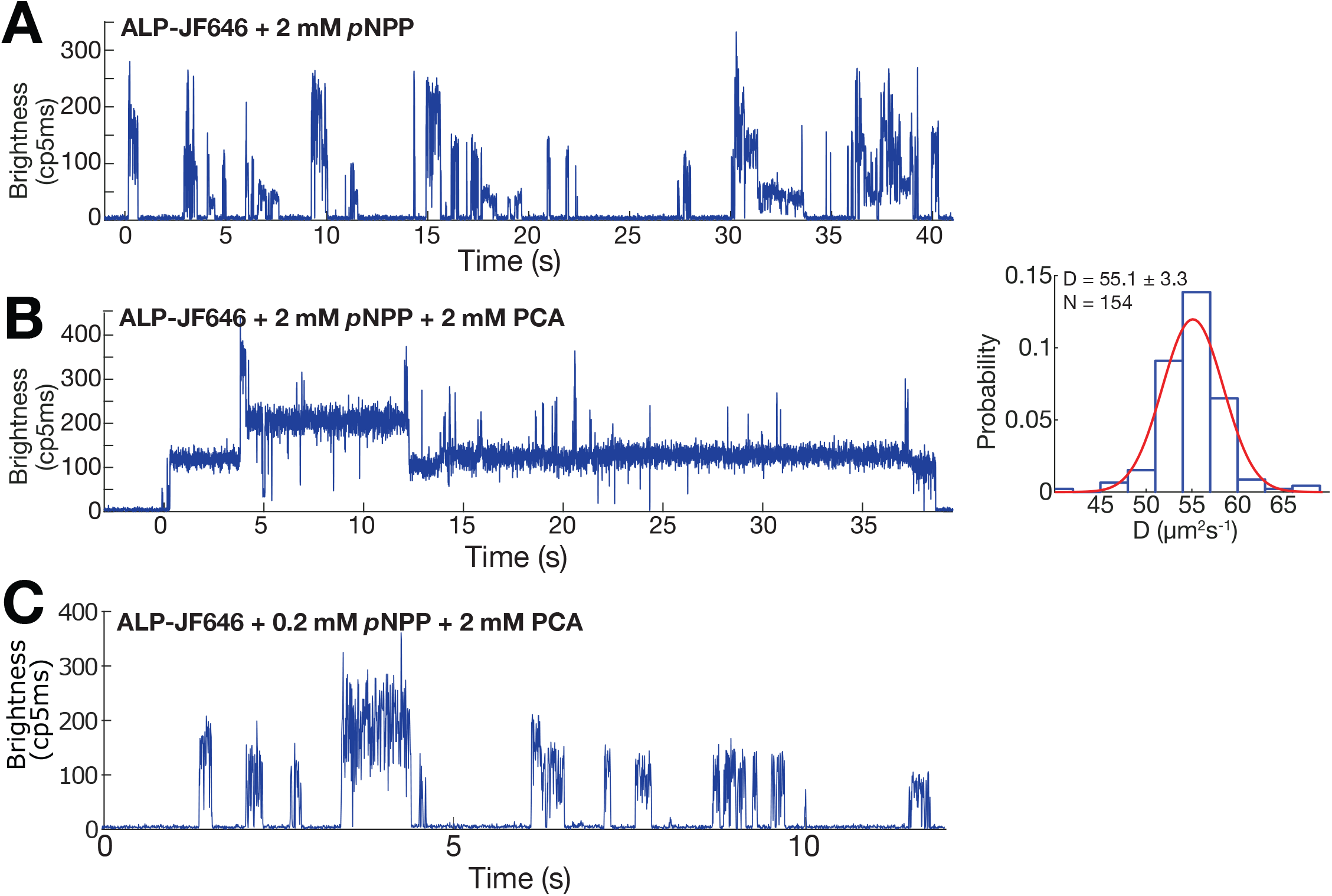
Additional ABEL trap data of ALP-JF646 under different buffer compositions. (A) A representative ABEL trap intensity trace of ALP-JF646 in the presence of 2 mM *p*NPP. Note the dye blinks rapidly. (B) Left, a representative ABEL trap intensity trace of ALP-JF646 in the presence of 2 mM *p*NPP and 2 mM PCA. PCA alone greatly suppresses *p*NPP-induced dye blinking. Right, D histogram of ALP under the trapping condition on the left. (C) A representative ABEL trap intensity trace of ALP-JF646 in the presence of 0.2 mM *p*NPP and 2 mM PCA. Rapid blinking was also observed and understood as originating from an imbalance of oxidizer and reductant.

**Figure S7.**
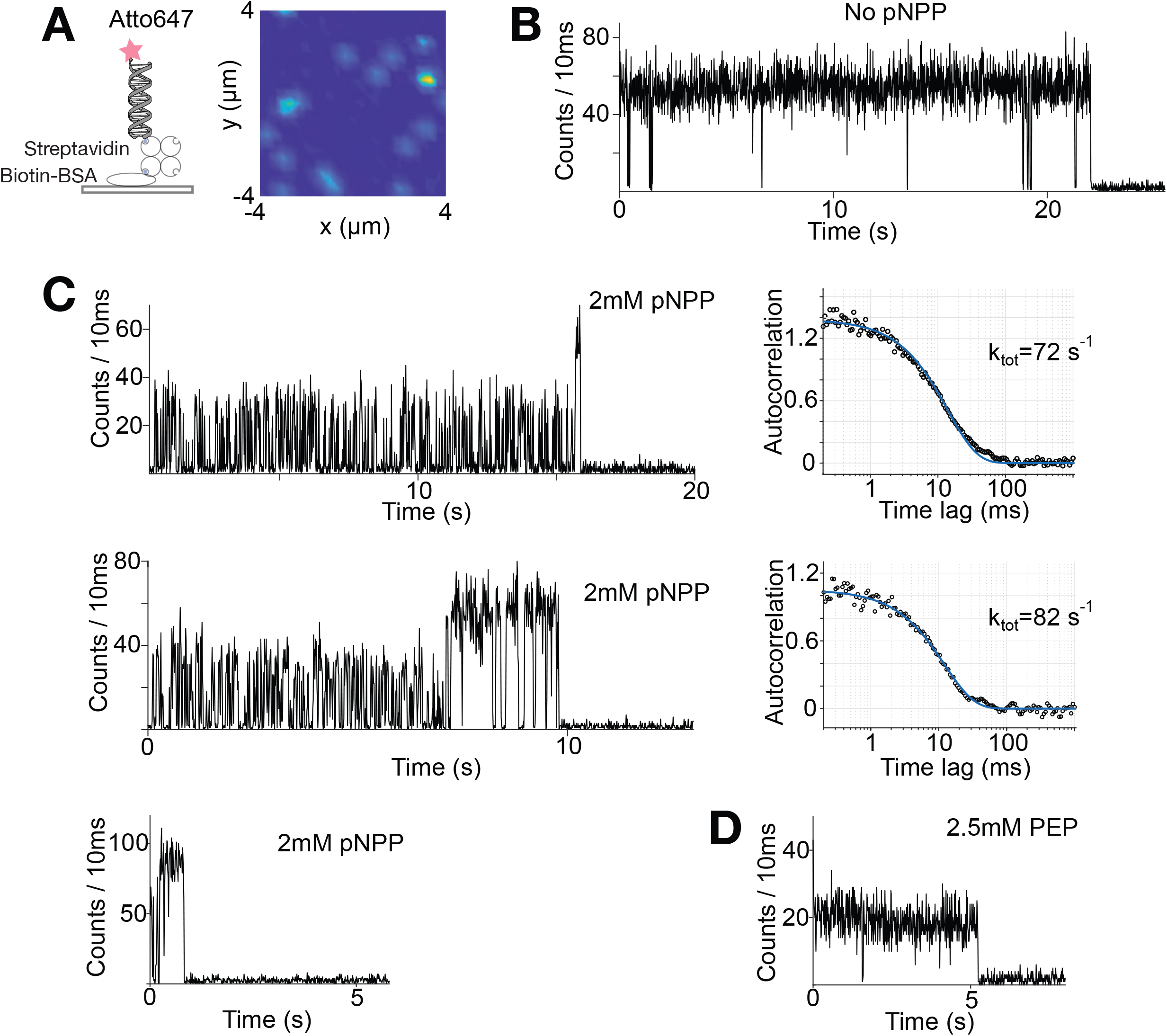
Surface-based single-molecule spectroscopy on Atto647-labeled double stranded DNA molecules. (A) Left, single Atto647-dsDNA molecules tethered to the surface via biotin-streptavidin interactions. Right, a representative image showing an 8μm ×8μm field of view, acquired in 1×PBS buffer without substrate. (B) A representative intensity (photon counts per 10 ms) trace of a single Atto647-dsDNA molecule without *p*NPP. (C) Representative intensity traces of single Atto647-dsDNA molecules in the presence of 2 mM *p*NPP. About 50% of the molecules show rapid on-off blinking. The blinking traces are analyzed by fitting the intensity autocorrelation function with a single-exponential decay. The extracted rates confirm the time scale of blinking to be ~ 10 ms. (D) A representative intensity traces of single Atto647-dsDNA molecule in the presence of 2.5 mM PEP. PEP does not induce dye blinking.

**Figure S8.**
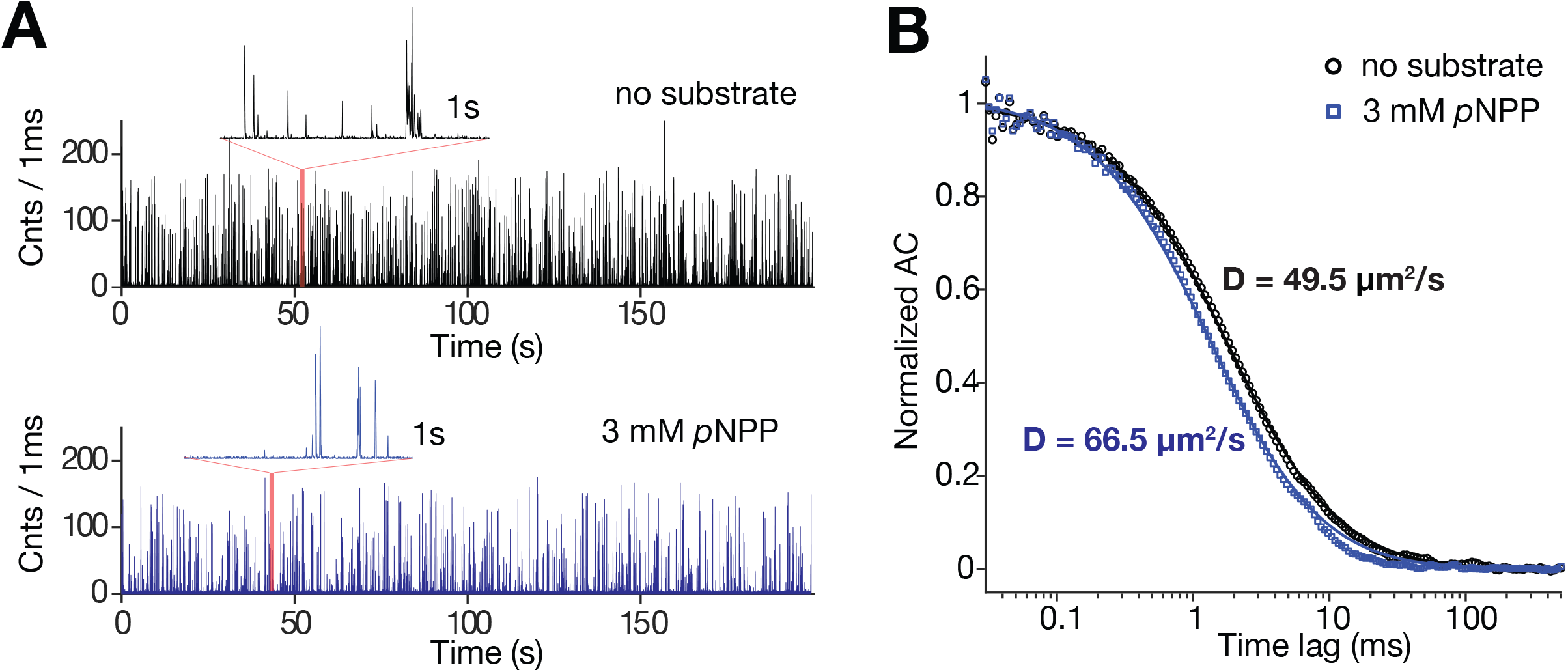
Monte-Carlo simulation of FCS experiments with substrate-induced quenching. (A) Representative intensity traces generated by the simulation (black: no substrate, blue: 3mM substrate). (B) Calculated intensity autocorrelation functions from (A) (symbols) and corresponding fits to a one-component simple diffusion model. The extracted apparent diffusion coefficient values are shown.

## Supplementary Information

### Materials and Methods

#### General Materials

All DNA modifying enzymes were purchased from New England Biolabs (NEB). Oligonucleotides were purchased from Integrated DNA Technology (IDT). Standard salts and buffer components were purchased from Sigma Aldrich. Cloning and DNA template construction follows standard molecular biology techniques unless otherwise noted.

#### Purification of ALP

Alkaline phosphatase (ALP) from bovine intestinal mucosa was purchased from Sigma-Aldrich (Cat # P7923). The proteins were dialyzed against 1 L of HN buffer (20 mM HEPES, pH 8.0, 100 mM NaCl) with 1 mM DTT, and then concentrated with the same buffer using Amicon Ultra-15 Centrifuge Unit with Ultracel-30 membrane (Millipore Sigma). Concentrated proteins were loaded on a Superdex 200 10/300 GL size exclusion column (GE Healthcare) in HN buffer with 1 mM DTT and eluted in 0.5 mL fractions. Pure fractions of the enzymes in correct oligomeric states were pooled, concentrated, aliquoted, flash frozen in liquid nitrogen and stored at −80 °C. Protein concentration was estimated from absorbance at 276 nm using extinction coefficient of 49070 M^−1^cm^−1^ (per monomer).

#### Cloning, expression and purification of sgPP

*Streptococcus gordonii* inorganic pyrophosphatase (*sg*PP) gene (UniprotKB-P95765) was synthesized as a gBlock gene fragment from IDT. The gBlock gene fragment also contains tags and adaptor sequences and was subcloned in a 1b vector (pET-His6-TEV LIC cloning vector, Addgene plasmid #29653) using NEBuilder HiFi DNA Assembly Cloning Kit (NEB cat#E5520S). The cloned *sg*PP gene plasmid (1b-*sg*PP) contains a N-terminal Ybbr tag (1) and C-terminal Sortag (2) followed by a hexa-Histidine tag. The 1b-*sg*PP-S195C mutant was cloned using Q5 Site-Directed Mutagenesis Kit (NEB cat#E0552S). The sequences of the forward and reverse oligos for cloning are:

*sg*PP S195C-F: 5’-GCGGGTACCAATCTTGCCTgcAAGTCTGCTGAGGAACTG-3’
*sg*PP S195C-R: 5’-TTTTAACATAGCCAGACCGTACTCCTCTAAATTGAC-3’

To express *sg*PP or *sg*PP-S195C, the plasmid harboring the gene was transformed into Rosetta™ 2 Competent Cells (Millpore Sigma Cat#71402-3). A single colony was inoculated into 10 mL of LB media supplemented with 25 μg/mL kanamycin and 50 μg/mL chloramphenicol and grown overnight at 37 °C. The cells were then diluted into 1 L of Terrific Broth (TB) medium supplemented with 1% glucose, 1 mM MgSO_4_, 25 μg/mL kanamycin and 50 μg/mL chloramphenicol and grown at 37 °C, 250 rpm until OD_600_ reaches ~0.6. Protein expression was induced with 0.1 mM IPTG and growth was continued at 17 °C for 16 h. Cells were harvested by centrifugation at 4000 rpm for 20 mins, resuspended in lysis buffer (20 mM Tris-Cl, pH 8.0, 500 mM NaCl, 20 mM imidazole, 10% glycerol, 0.1 mg/mL lysozyme, supplemented with 1 tablet of EDTA-free proteinase inhibitor), lysed by sonication (10 s pulse, 10 s pause for a total of 5 mins on ice bath), and clarified by centrifugation at 18,000g for 30 mins. Clear lysate was filtered with 0.22 μm syringe filter and loaded on a 5 mL HisTrap HP column (GE Healthcare Cat#17524801). The column was washed with 25 mL HisP-buffer (20 mM Tris-Cl, pH 8.0, 500 mM NaCl) containing 50 mM imidazole, and eluted with 50 mL of HisP-buffer containing 50-300 mM imidazole gradient. The eluted fractions were loaded on a 12% SDS-PAGE gel and stained with AcquaStain Protein Gel Stain (Bulldog Bio Cat#AS001000). Fractions with abundant target protein were pooled and dialyzed against 1 L of HN buffer at 4 °C overnight. The protein was concentrated by using Amicon Ultra-15 Centrifuge Unit with Ultracel-30 membrane and loaded on a Superdex 200 10/300 GL size exclusion column (GE Healthcare) in HE buffer and eluted in 0.5 mL fractions. Pure fractions of the enzymes in correct oligomeric states (dimer) were pooled, concentrated, aliquoted, flash frozen in liquid nitrogen and stored at −80 °C. Protein concentrations were estimated from absorbance at 276 nm using extinction coefficient of 14900 M^−1^cm^−1^ for *sg*PP or *sg*PP-S195C (per monomer).

#### Labeling of ALP, TIM and sgPP-S195C

ALP was labeled using JF 646-NHS (*N*-hydroxy-succinimide) at a protein (dimer): dye molar ratio of 1: 5 in 0.1 M sodium bicarbonate buffer (pH 8.3-8.5) at room temperature for 2 hours. Excess dyes were removed by using 5 mL Zeba Spin Desalting Columns, 40K MWCO (Fisher Scientific Cat#87771) and buffer exchanged in HN buffer. The desalting step was repeated to ensure as complete as possible of free dye removal. The absorption spectrum (230-700 nm) of the labeled protein was measured on a NanoDrop 2000 Spectrophotometer. Protein and dye concentrations were estimated using the extinction coefficient of the protein (49070 M^−1^cm^−1^ per monomer at 276 nm) and the dye (152,000 M^−1^cm^−1^ per dye at 646 nm), assuming an A280 correction factor of 0.19. Labeling of ALP with Atto647N follows similar procedure. TIM was purified and labeled as previously described (3).

*sg*PP-S195C was labeled using Atto647N maleimide (Sigma-Aldrich Cat#05316) at a protein: dye molar ratio of 1: 10 in labeling buffer (50 mM Tris-Cl, pH 7.0, 100 mM NaCl) at room temperature for 2-4 hours. Excess dyes removal, absorption measurement and protein/dye concentration estimation were similar as described above. The extinction coefficient and A280 correction factor for Atto647N-maleimide were 152,000 M^−1^cm^−1^ per dye at 644 nm, and 0.05, respectively.

#### ALP enzyme activity assays

For ALP activity with *p*NPP, ALP was diluted to 0.05 nM in 20 mM Tris, 100 mM NaCl, 1 mM MgCl_2_, 20 μM ZnCl_2_, pH 10.0. 90 μl of the ALP solution was transferred to a 96 well plate, and 10 μl of 10x concentrated *p*NPP was added and the absorbance at 405 nm was monitored (extinction coefficient = 18000 M^−1^cm^−1^) at room temperature for 3 minutes.

Activity assays were performed to compare the activity of bALP with *p*NPP in the presence and absence of Trolox and PCA/PCD. Under both conditions the reaction buffer was composed of 25 mM HEPES pH 8.0, 100 mM NaCl, 10 mM MgCl_2_, 2 mM *p*NPP, and bALP-JF646 at nanomolar concentration. The assay with Trolox and PCA/PCD additionally contained those reagents at the concentrations: ~3 mM Trolox, ~50 nM PCD, and ~3 mM PCA. The absorbance at 405 nm was monitored at room temperature over 5 minutes using a Beckman DU650 Spectrophotometer.

For ALP activity with phosphoenolpyruvate (PEP) or adenosine monophosphate (AMP), ALP was diluted to 1 nM in 20 mM Tris, 100 mM NaCl, 1 mM MgCl_2_, 20 μM ZnCl_2_, pH 8.0. 10 μl of 10x substrate was added to 90 μl ALP solution at various time points (30, 60 and 120 sec for PEP and 120, 240 and 420 sec for AMP). Final substrate concentrations are 0.5 and 1 mM for PEP and 0.625 and 1.25 mM for AMP. 10 μl of the enzyme reactions were then diluted to 100 μl with Tausky-Shorr Reagent (1% Ammonium molybdate, 1N Sulfuric Acid, 5% Ferrous Sulfate). Serial dilutions of a phosphate standard (Sigma-Aldrich P3869) was used to create a standard curve to calculate the phosphate concentrations in the enzymatic reactions.

#### FCS measurements

FCS was performed on a custom-built confocal microscope. A 633nm laser line (JDSU 1135P HeNe) was focused into the sample using an infinity-corrected oil immersion objective (UApo N 100X NA=1.49, Olympus), to give a peak intensity at the focus of ~1 kWcm^−2^. Emitted fluorescent photons were collected by the same objective, focused onto a 100μm pinhole, collimated and then refocused onto the active area of an avalanche photodiode (APD) detector (SPCM-AQRH-24-TR). The APD output was connected to an FPGA board which records photon arrival times with 8.3 ns resolution.

The sample chambers were made by bonding a plasma cleaned 1.5H glass coverslip to a silicone hybridization chamber (Grace Bio-labs SecureSeal). Experiments were performed at a typical volume of 20 μL and a labeled enzyme concentration of around 0.5 − 2.5 nM. All recordings were made ~ 15 μm above the coverslip surface to avoid surface effects.

Intensity autocorrelation curves were calculated using custom written software in MATLAB and C++ using a flexible binning algorithm (4). The calculated autocorrelation curves were fitted to extract the diffusion coefficient by a model assuming a single diffusing species:

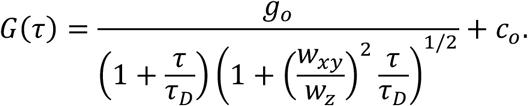

The amplitude *g*_*o*_;, the diffusion time *τ*_*D*_, and a constant offset *c*_*o*_ were the fitting parameters. *w*_*xy*_ and *w*_*z*_ represent the transversal and longitudinal dimensions (1/e^2^ radii) of the focal volume respectively and were measured before the experiment by scanning a 200 nm fluorescent bead (Invitrogen Cat. No. F8806) to map out the focal volume (5). The diffusion coefficient was calculated using 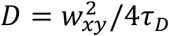.

#### ABEL trap based single-molecule diffusometry

The ABEL trap hardware was implemented as previously described (6, 7). Briefly, we used fast laser scanning and photon-by-photon mapping to sense the position of a single molecule in real time. The raw measurements were refined using a hardware-implemented Kalman filter. Based on the refined position estimates, feedback voltages were calculated and applied to a microfluidic sample holder to counteract Brownian motion. In this work, excitation was provided by a 637nm laser (Coherent Obis) with an average intensity of ~600 W/cm^2^ at the sample plane. The fluorescence signal was collected by a silicone immersion objective (100X, NA1.35, Olympus) and focused onto an avalanche photo detector (SPCM-AQRH-24-TR) using a confocal geometry. The detection area at the sample plane was restricted to 4μm in diameter using a 400μm (Thorlabs) pinhole.

The microfluidic sample holder was made of UV-grade fused silica and fabricated in Princeton University’s nanofabrication facility using a modified version of the published recipes (8). The dimensions of the trapping region are approximately 3 μm × 3 μm × 0.7 μm, with the smallest dimension (0.7 μm) set by the depth of the microfluidic chip. Before measurements of protein sample, the microfluidic chip was passivated with polyethylene glycol (PEG). This was done by first cleaning the chip using Piranha solution (3:1 mixture of H_2_SO_4_ and hydrogen peroxide), activating the surface silanol groups using 1M potassium hydroxide followed by a 24 h incubation with methoxy-PEG-silane (Gelest SIM6492.73-1GM).

The diffusion coefficient of individual trapped molecules was extracted using a maximum-likelihood framework as previously described (7). Briefly, the algorithm uses the photon-stamped position sequence of a single trapped molecule and the feedback voltages as inputs and utilizes an iterative procedure to reconstruct the molecule’s motion trajectory while being trapped. This motion can then be decomposed into a voltage-dependent component and a voltage independent, stochastic component which yields the diffusion coefficient. In this work, diffusion coefficient is estimated in 100ms time windows and averaged to yield a single molecule’s mean D value.

#### Surface-immobilized single-molecule measurements

Single-molecule spectroscopy of JF646 and Atto647 was carried out using surface immobilized dye-labeled DNA duplexes on the same custom-built confocal microscope used for FCS measurements. To generate dye-labeled DNA samples, amino-modified single-stranded oligonucleotides (45 bases ssDNA, 5’-Amino-AGCTGGATCCTAATACGACTCACTATAGGGAGACCACAACGGTTT-3’) were labeled with JF646 or Atto647 using standard NHS-ester chemistry. Specifically, 20 μM of amino-modified ssDNA was incubated with ~200 μM dye-NHS in 0.2M sodium bicarbonate buffer (pH=8.6) for > 4 hours, protected from light. Labeled ssDNA were purified by P6 spin column (BioRad) and hybridized with its biotinylated complementary strand (5’-Biotin-AAACCGTTGTGGTCTCCCTATAGTGAGTCGTATTAGGATCC AGC T-3’). To immobilize the biotinylated, dye-labeled duplex, a glass coverslip (Zeiss 1.5H) was first plasma cleaned (air, 600 mTorr, 5 min) and incubated in 2 mg/ml biotinylated BSA (Pierce 29130) solution for 10 minutes. After rinsing 3 times with PBS, the surface was incubated with streptavidin (Pierce 21122, 0.1mg/ml) for 5 minutes and rinsed again 3 times with PBS. Then, the biotinylated, dye-labeled duplex was incubated on the surface for 2 minutes. Before imaging, the surface was rinsed 3 times and covered with imaging buffer (1X PBS, pH 7.4 for Atto647 and 20 mM HEPES, 100 mM NaCl pH 8.0 for JF646).

To record single-molecule emission traces, the sample was first raster scanned using a piezo stage (P-562.3CD) to identify isolated spots of individual molecules (Figure S3A and S7A, typically 8 μm×8 μm, with 0.2 μm step size and 20 ms dwell time). The molecule of interest was then moved to the excitation laser spot (peak intensity ~300 W/cm^2^) to record emission intensity as a function of time. All steps were controlled by custom written software in LabView.

#### Single-particle Tracking (SPT) experiments and analysis

SPT acquisition was realized following Ref. (9). Briefly, single-molecule imaging was performed on a custom-built Nikon TI microscope (Nikon Instruments Inc., Melville, NY) equipped with a 100x/NA 1.49 oil-immersion TIRF objective (Nikon apochromat CFI Apo TIRF 100x Oil), EM-CCD camera (Andor, Concord, MA, iXon Ultra 897), a perfect focusing system to correct for axial drift and motorized laser illumination (Ti-TIRF, Nikon), which allows an incident angle adjustment to achieve highly inclined and laminated optical sheet illumination (10). Excitation was achieved using the following laser line: 633 nm (1W, Genesis Coherent, Pala Alto, CA). The excitation lasers were modulated by an acousto-optic tunable filter (AA Opto-Electronic, France, AOTFnC-VIS-TN) and triggered with the camera TTL exposure output signal. The laser light is coupled into the microscope by an optical fiber, reflected using a multi-band dichroic beamsplitter (405 nm/488 nm/561 nm/633 nm quad-band, Semrock, Rochester, NY) and then focused in the back focal plane of the objective. Fluorescence emission light was filtered using a single band-pass filter placed in front of the camera (Semrock 676/37). The microscope, cameras, and hardware were controlled through NIS-Elements software (Nikon).

The framerate was set at 1.7ms/frame (~590 Hz, consisting of a ~1.3 ms exposure time followed by a ~0.447 ms camera dead time). The main excitation laser (633 nm) was pulsed for 1 ms starting at the beginning for the 1.7 ms camera exposure time. The camera settings were as follows: frame transfer mode; vertical shift speed: 0.9 ms; ROI: height 22 pixels, width 512 pixels. Each condition was imaged for more than 250,000 frames corresponding to ~7 min. The enzyme concentration was optimized to keep an average molecule density of < 0.5 localizations per frame. Maintaining a very low density of molecules is necessary to avoid tracking errors. The main excitation laser was used at maximal power. The results presented here are the results of independent measurements performed at least twice on different days.

To extract diffusion coefficients, SPT data was processed following a published protocol (9). SPT data was analyzed (localization and tracking) and converted into trajectories using a custom-written Matlab implementation of the multiple-target tracing algorithm (11) (available at https://gitlab.com/tjian-darzacq-lab/SPT_LocAndTrack) with the following settings: Localization error: 10-6.25 ; deflation loops: 0; Blinking (frames): 1; max competitors: 3; max D (μm^2^/s): 120. The SPT trajectory data was then analyzed using the Python version of Spot-On (https://gitlab.com/tjian-darzacq-lab/Spot-On-cli, version 15.10) and the following parameters: dZ = 0.7 μm; GapsAllowed = 1; Time-Points: 4, useEntireTrajectory = yes; NumberOfStates = 2; FitLocError = no; LocError = 0.035 mm; D_Free_2State=[0.15;50]; Frac_Bound = [0, 0.001] (to constrain the fitting to only one state).

To compare the brightness and density of the detected spots, two movies (one without *p*NPP, the other with *p*NPP) acquired on the same day at less than one hour interval with nearly-identical microscope settings (including TIRF angle) were used. Firstly, the movies were cropped to identical lengths (50,000 frames) and the mean pixel value across the whole movies was computed (321.0 ±101.7 without *p*NPP vs. 288.9 ±88.3 with *p*NPP, ns), confirming that the imaging conditions were indeed similar. Second, the movies were processed with identical parameters using TrackMate (12). Parameters: Spot size: 0.48 μm ; Use median filter : checked ; Threshold: 200 (pixel intensity). The results were then exported in a CSV table and both the number of detections and the ‘MEAN_INTENSITY’ column were analyzed.

#### Monte-Carlo simulation of FCS experiments

Simulation was conducted by keeping track of the 3D position and photon emission of two molecules diffusing in and out of a Gaussian-shaped focal volume with w_xy_=0.6 μm and w_z_=2 μm. Molecules were restricted to diffuse inside a box with dimensions of 2.4μm×2.4μm×7μm with reentrant boundary conditions. The Brownian motion was simulated by drawing Gaussian-distributed position increments 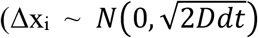, 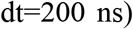 of all three dimensions. The photon emission process of one molecule was simulated by drawing Poisson random variables with mean 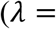 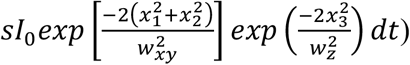 determined by the intrinsic brightness I_0_ (200 cnts/ms) coordinates (x_1-3_) with respect to the Gaussian focal volume and emission states (s), which in this work, toggles between s=1 (on state) and s=0 (off state) with a pair of rates k_1_ and k_2_. The total emission rate was the sum of two molecules and a constant background rate (0.8 cnts/ms). The simulated photon arrival time trajectory was then subject to the same analysis pipeline as in the FCS experiments to compute and fit the intensity autocorrelation function. For each condition, the simulation was run for 1×10^9^ time steps (200 seconds in total).

